# Discovery & Evaluation of novel fluorescence molecules for selective recognition of G-quadruplexes structure

**DOI:** 10.1101/2023.07.08.548211

**Authors:** Neha, Prashant Ranjan, Parimal Das, Surendra Kumar, Roop Shikha Singh, Daya Shankar Pandey

## Abstract

Currently, G-quadruplex structure targeting strategies are considered as a promising anticancer approach. In the search of selective and potent G-quadruplex binders, Here we discuss an analysis of a few chroman derivatives ligands: (A) chroman 7-[2-pyrrolo]-pyrrole-[1,2-a]12H pyrrolino[2,3-b]chroman-4-one, and (C) 4-methyl-7-[2-pyrrolo]-pyrrole[1,2-a]12H pyrrolino[2,3-b]chroman-4-one and their respective borondifluoride complexes B and D as a quadruplex targeting compounds which found to stabilize G-quadruplex structure. To investigate the binding characteristics of these molecules with G-quadruplex vs. duplex selectivity, *In vitro* biophysical studies were performed by steady-state fluorescence, UV-visible titration, fluorescent TO displacement assay, CD thermal melting, circular dichroism spectroscopy, and cellular imaging by employing both telomeric and PRCC G-quadruplex forming sequences. Our investigation shows that these chromam ligands and their complexes are able to selectively bind and stabilize parallel and mixed hybrid topology of G-quadruplex both *In vitro* and in cellular conditions. A molecular docking study also suggests the binding of these compounds with G-quadruplex conformation. Collectively our study suggests these chroman complexes as a potentially useful fluorescent chemical product for G-quadruplex specific ligands and expands an option for G-quadruplex targeting ligands.

## 1. Introduction

In guanine-rich sections of the genome, nucleic acids have a distinctive secondary structure called the G-quadruplex. Over the past two decades, G-quadruplexes have become more well-known for their function in the regulation of gene activity, the management of anti-tumor activity, and aging. Numerous reviews have been done on the physiological importance and relevance of these non-canonical structures in relation to cancer (Verma & Das, 2018b, 2018a). G-quadruplexes are thought to play roles in cancer pathogenesis and prognosis, which has prompted researchers to look for small molecule ligands that might bind to and influence the activity of these structures. In similar a direction, G-quadruplex structures may now be recognized specifically using small molecule fluorescent probes. These have made it possible to directly visualize and track these structures (Chilka et al., 2019; Sun et al., 2019). These structures are now considered to be worthwhile targets in molecular studies thanks to recent and conclusive evidence of the synthesis of DNA and RNA G-quadruplexes in cells (Varshney et al., 2020)(Das & Verma, 2020)(Das & Verma, 2020)(Das & Verma, 2020). The necessity of creating multi-tasking ligands with appealing quadruplex interacting properties (affinity, structural selectivity), as well as additional features that make them usable for detecting quadruplexes, for better understanding the effects of their stabilization by external agents, is made clear by G-quadruplex targeting strategies. The majority of the G-quadruplex ligands identified thus far directly derive from the duplex-DNA intercalator pool’s fine structural tunings (Ohnmacht & Neidle, 2014) (Vy Thi Le et al., 2012). This strategy seems a little contradictory given that the main undesirable event for a chemical to be a viable ligand is interaction with duplex-DNA. Ingenious structural modifications, such as enlarging the aromatic core or increasing the side arms, were cleverly used to reduce interactions with duplex binding sites (base pairs, grooves) in favor of the quadruplex one (external G-quartet) (Haider et al., 2011), but in most cases, the selectivity level achieved still remains debatable, especially given that the vast majority of DNA is found in cell nuclei in its duplex form. A sensible and effective technique is the discovery of fluorescent ligands that target or bind G-quadruplexes. For this two fluorescent ligands and their complexes were evaluated for their G-quadruplex interacting property by a series of biophysical approaches. Due to their widespread application in numerous fields, dipyrromethenes (also known as dipyrrins) have attracted a great deal of interest among scientists (Falk, 1989; Harmjanz et al., 2000; Lindsey, 2010; Lindsey et al., 1994). The planar dipyrrin unit functions as a flexible ligand and holds a respectable place in coordination chemistry thanks to the presence of two nitrogen donor atoms. These complexes attract attention for their DNA binding behavior due to the photophysical and lasing characteristics of borondifluoride dipyrrinato complexes (BDPs), as well as their superior photostability, flexible chemistry, and prospective applications in diverse fields (Bura et al., 2011; Treibs & Kreuzer, 1968; Ulrich et al., 2008; Welch, 1999) Therefore, it has been interesting to investigate the DNA binding properties of BDP derivatives and their analogues, which have relatively high fluorescence quantum yields and substantial Stokes shifts (SS). Furthermore, chromone derivatives are abundant in nature and display a variety of biological and pharmacological properties. These are present in all green plant cells and are anticipated to take part in photosynthetic processes. Chromones can be used in photo-induced reactions since they are also photoactive and can produce a wide range of heterocyclic compounds (Mukohata et al., 1978; Singh et al., 2013). In the present study few chroman derivatives ligands: chroman 7-[2-pyrrolo]-pyrrole-[1,2-a]12H pyrrolino[2,3-b]chroman-4-one (A), and 4-methyl-7-[2-pyrrolo]-pyrrole[1,2-a]12H pyrrolino[2,3-b]chroman-4-one (C) and their borondifluoride complexes: complex of chroman derivative ligand A is B, and of C is D (Figure 1) which exhibit unusual structure with relatively high fluorescent quantum yield and having good thermal stabilities, and emissive properties (Singh et al., 2013) were subjected to investigate their G-quadruplex binding capability.

**Figure 1:**
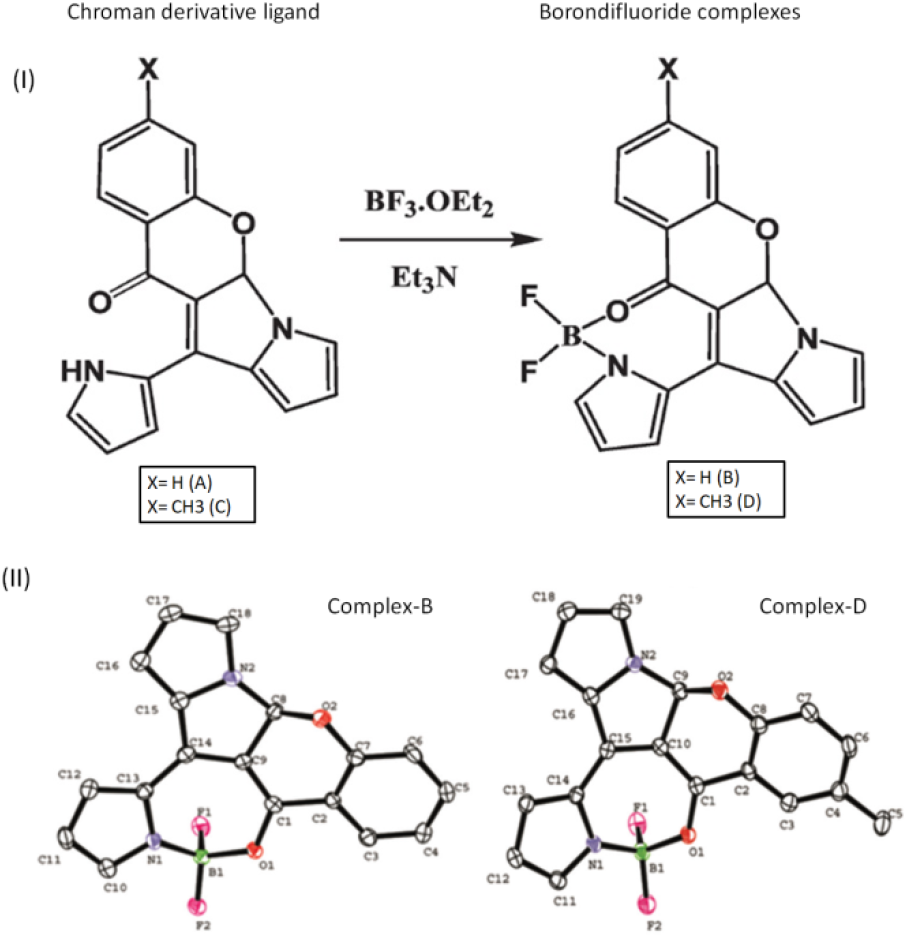
Schematic presentation of chroman ligands and their borondifluoride complexes (I). Molecular structure presentation by Oak Ridge Thermal Ellipsoid Plot view of complex B and D (II). Adapted from (Singh et al., 2013)

## 2. Materials and Methods

### 2.1 Materials and Reagents

The HPSF grade 22 bp human telomeric sequences (TelQ), 27 bp PRCC DNA oligonucleotides (PRG4Q) bearing G-quadruplex forming sequence and its mutated version PRG4M (where repetitive guanine bases get altered with adenine base) were obtained from Eurofins Genomics India Pvt. Ltd. and utilized as received. We utilized salmon sperm DNA as the duplex or B-DNA, which was obtained from Sigma-Aldrich and used without additional purification. Thioflavin-T was also bought from Sigma-Aldrich. Chroman derivatives ligands and their borondifluoride complexes were provided in powdered form by Dr. Roop Shikha Singh, Assistant Professor, Department of Chemistry, Banaras Hindu University, Varanasi, India. Both ligands (A and C) were dissolved in MQ water while both complexes (B and D) were dissolved in DMSO to make stock solution. Fetal bovine serum (FBS), penicillin-streptomycin antibiotic cocktail, Dulbecco’s Modified Eagle’s Medium (DMEM) medium, and MTT (3-(4,5-dimethylthiazol-2-yl)-2,5-diphenyltetrazolium bromide) were all bought from Himedia.

### 2.2 Sample Preparation

The samples of oligonucleotides were dissolved and heated in 10 mM Tris at pH 7.4 and 100 mM KCl at 95°C for 5 min in a heat block followed by gradual cooling at room temperature for overnight to anneal into G-quadruplex conformation and stored at 4°C. Following oligonucleotides were utilized in this study.

i. PRCC (PRG4Q): 5’-GTTGGGGAGGGACTGGGATTGGGGTTG-3’
ii. Mutated PRCC (G4M): 5’-GTTGAGGAGAGACTGAGATTGAGGTTG-3’
iii. Human Telomeric (TelQ): 5’-AGGGTTAGGGTTAGGGTTAGGG-3’

### 2.3 Circular Dichroism Spectroscopy

To capture the formation of the G-quadruplex structure, JASCO J-1500 CD Spectrophotometer has been used. CD measurements were performed in a quartz cuvette with 2.0 mm route lengths at 25°C at a wavelength from 220 to 320 nm. Bandwidth and data pitch were set to 1.0 nm and 0.2 nm respectively. The data interval time was 1 sec. The CD spectra were recorded at averages across three scans at a speed of 50 nm min^-1^, with the required baseline correction applied from the blank buffer of 10 mM Tris at pH 7.4 with 100 mM KCl.

### 2.4 UV-Visible titration

The UV-titration was undertaken with UV-1800 UV/Visible Scanning Spectrophotometer using a quartz cuvette with 1 mm path lengths. The buffer solution used for all titration tests contained 10 mM Tris, and 100 mM KCl, at a pH of 7.4. A cuvette containing 30 µM of compound A to D was subjected to a progressive addition of PRCC G4Q and telomeric G-quadruplex solution (1-10 μM) to perform a UV-Vis absorption titration. At room temperature, absorption spectra between 200 to 800 nm were scanned. The titration was completed when, after three repeated additions of tetrad oligos, neither the wavelength nor the intensity of the compound’s absorption band shifted.

### 2.5 Thermal Melting Study

At a heating rate of 1°C min^-1^, DNA melting was carried out both with (10 μM) and without the molecules using a CD spectropolarimeter (JASCO J-1500) equipped with a Peltier temperature controller. 5 µM of oligo samples were annealed to achieve a G-quadruplex structure as mentioned in the above sample preparation section. All melting studies were monitored by heating the samples from 30°C to 90°C at 260 nm at a speed of 1°C min^-1^. The formula used to determine the fraction folded was Qt-Qu/Qf-Qu, where Qt represents the molar ellipticity at temperature T and Qu and Qf represent the molar ellipticities of the fully unfolded (highest temperature) and fully folded (lowest temperature), respectively.

### 2.6 Fluorescence Spectroscopy

A Perkin Elmer LS55 fluorescence spectrometer was used to record the fluorescence spectra at a temperature of 25°C. All tests were carried out using a quartz cuvette with a 1 mm path length. Excitation and emission slits for fluorescence measurements were 10 and 7 nm respectively, and the scanning speed was adjusted at 500 nm/min. To determine the binding constants, each compound was titrated with folded DNA G-quadruplex samples: PRG4Q and TelQ. The titration studies involved raising the DNA G-quadruplex concentrations from 1 to 10 μM, which was added to the fixed concentration of 30 µM of a compound with 5 min incubation at room temperature. Each compound was excited at its corresponding excitation wavelength (mentioned in Table 1).

**Table 1:** Absorption/Emission maxima of compound A-D

| Compounds | UV abs $\lambda/\text{nm}$ | Fluo $\lambda/\text{nm}$ | |
| --- | --- | --- | --- |
| | | $\lambda_{\text{ex}}/\text{nm}$ | $\lambda_{\text{em}}/\text{nm}$ |
| A | 413, 463 | 463 | 520 |
| B | 408, 462, 490 | 462 | 519 |
| C | 481 | 481 | 516 |
| D | 482 | 482 | 521 |

### 2.7 Fluorescence Intercalating Displacement Assay

Fluorescence intercalator displacement titrations were conducted in the appropriate buffers at room temperature. First DNA samples were folded into G-quadruplex conformation as mentioned in the sample preparation section. The DNA solutions were made in a 2:1 ratio by combining 10 µM of Thiazole Orange (TO) with 5 µM of G-quadruplex folded oligonucleotides. Fluorescence spectra were recorded with an excitation wavelength of 509 nm and emission wavelength of 400-700 nm at 25°C with increasing quantity (0.25-30 μM) of each of the compounds sequentially after a 5 min equilibration time. Excitation and emission slits for fluorescence measurements were 10 and 5 nm respectively and the scanning speed was adjusted at 500 nm/min.

### 2.8 Cell culture and immunofluorescence

In the 6 well plates, HEK293T cells were grown on methanol-charged cover slips. The following day, cells were exposed to 10 μM of compounds A - D and incubated for 24 hours in order to induce and stabilize the G-quadruplex structure in the cells’ genes. After 24 hours, cells were first washed with PBS, then fixed for 15 minutes with 3.7% paraformaldehyde, and then rinsed with PBS. After permeabilizing the cells for 30 min at room temperature with 0.1% Triton X-100, the cells were washed in PBS. 4% bovine serum albumin was used for blocking, which was done for two hours at room temperature. After blocking, cells were exposed to the G-quadruplex-specific antibody BG4 at 1:100 dilutions for 3 hours at room temperature in a moist environment. Next, the cells were washed with PBS, and the primary antibody (Flag) was added at 1:500 dilutions for another 3 hours of incubation at room temperature. Cells were washed with PBS after Flag antibody incubation, and then incubated for 3 hours at room temperature with secondary antibody Alexa Fluor-546 at 1:500 dilutions followed by PBS washing. Nuclear staining was carried out on cells using 1 g/ml DAPI for 15 min, followed by PBS washing and air drying. Slides were sealed with nail polish after being DPX mounted. Images were taken using a confocal microscope (ZWISS LSM 780). ImageJ software was used to analyze images or measure fluorescence intensity.

### 2.9 MTT assay

MTT assay was performed according to Neha *et al.,* 2023 (Neha et al., 2023). In brief DMEM-cultured HEK293T (Human Embryonic Kidney) cell line was cultivated in a humid incubator at 5% (v/v) CO2 with 10% fetal bovine serum (FBS), 100 g/ml streptomycin, and100U/ml penicillin antibiotic cocktail. Cells were cultivated in 96 wells plate for 24 hours at a density of 2×10^4^. The following day, cells were exposed to varying concentrations (5-50 μM) of each test compound for 24 hours. Following a 24-hour period, the media were changed to DMEM containing 0.5 mg/ml MTT and incubated at 37°C. After 4 hours, the MTT solution was removed, 100 μl of DMSO was added, and the mixture was then incubated in the dark for 10 minutes to solubilize the blue/purple color formazan. By measuring absorbance at 570 nm in a microplate reader (Bio-Rad), the final purple color was quantified, and the IC50 dose was determined using a linear regression curve.

### 2.10 Molecular docking study

The target crystal structure of parallel (1KF1), and mixed hybrid type human telomeric G-quadruplexes (2HY9) have been retrieved from the Protein Data Bank database (Berman et al., 2007). The 3D structures thus generated were visualized by Discovery Studio 4.0 (Studio, 2008) followed by the cleaning of structures such as water molecules from them. All four compounds’ structures were drawn using ChemSketch (Li et al., 2004), which was then used to transform the drawings into mol format. Open Babel (O’Boyle et al., 2011) further converts the mol file into the pdb format for docking experiments. In order to conduct a docking examination using compounds A to D, the crystal structures of parallel G-quadruplex and mixed hybrid G-quadruplex were employed as templates. Compounds were docked with selected targets using PatchDock Server (Schneidman-Duhovny et al., 2005) (parameter RMSD value 1.0 and complex type protein-small ligand). Then, the non-covalent interaction of docked complexes was visualized using PLIP (Salentin et al., 2015).

## 3. Result

### 3.1 Circular Dichroism

First, CD spectra were collected to confirm that both oligo samples were folded in G-quadruplex. There are two peaks in the CD spectra of PRCC G4Q that are indicative of parallel-stranded G4Q topologies: a negative peak at 240 nm and a positive peak around 263 nm. The CD spectra of human telomere G-quadruplex DNA, on the other hand, exhibited a hybrid-type mixed topology with a shoulder peak at 265 nm, a negative peak at 240 nm, and a positive peak at 290 nm (Figure 2). The parallel and mixed hybrid topology of PRG4Q and TelQ respectively were still maintained after the incubation with each individual compound suggesting the preservation of each G4Q architecture in the presence of compounds, however, peak intensity gets changed in both G4Q suggesting the interaction of each compound with G4Q structure (Figure 3).

**Figure 2:**
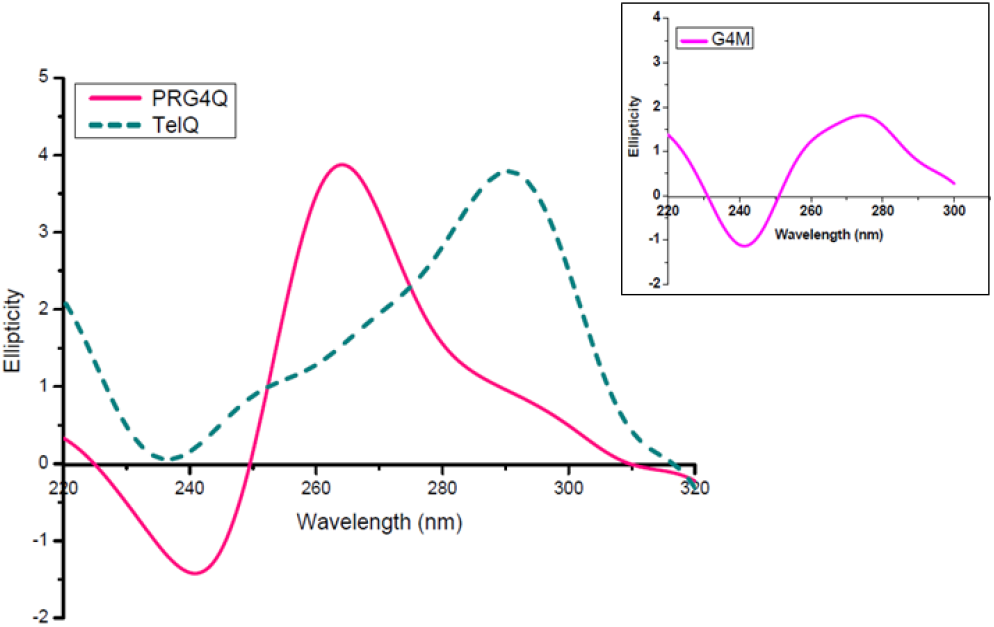
CD Spectra of PRG4Q (G-rich sequence within the PRCC stretch of *PRCC-TFE3* fusion gene) and human telomeric (TelQ) sequence. Showing formation of parallel G-quadruplex structure in PRG4Q and mixed hybrid G-quadruplex in TelQ. Spectra of the PRCC G4M sequence shows non G-quadruplex structure in inset

**Figure 3:**
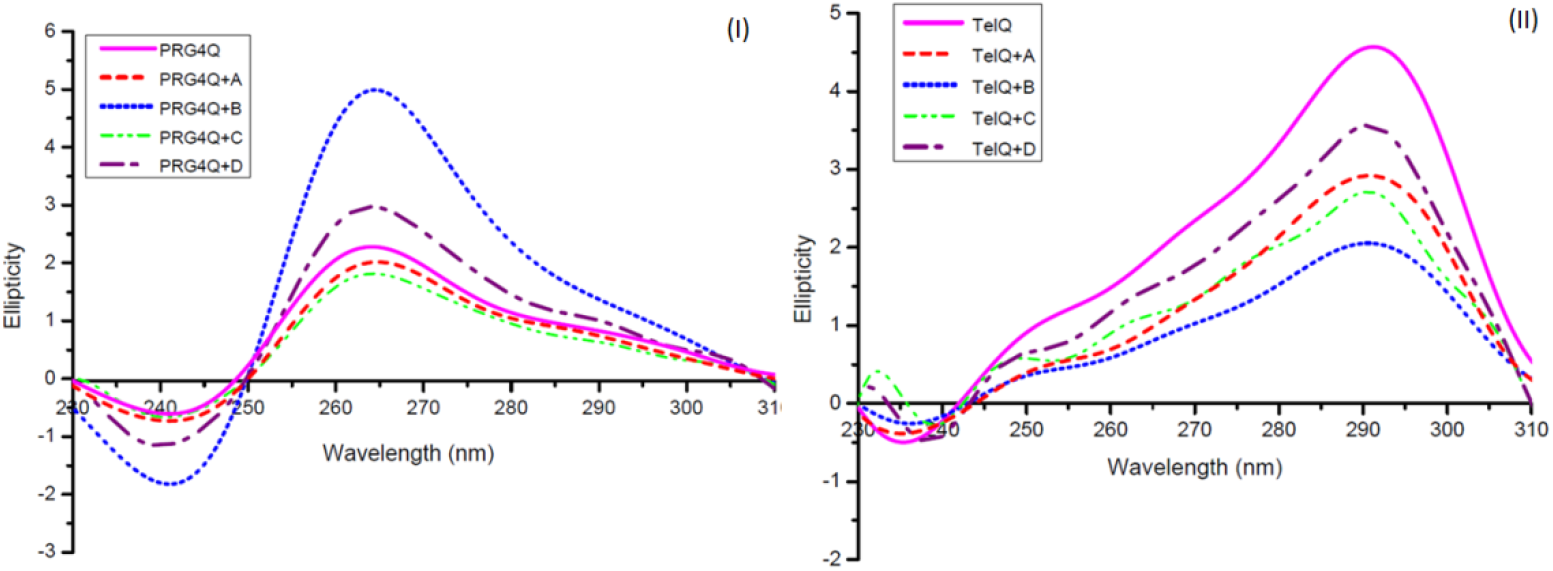
CD spectra of PRG4Q and TelQ in the absence or presence of 10 μM of compounds A to D. (I) Showing formation of parallel G-quadruplex structure in PRG4Q in the presence of compounds A-D. The peak intensity around 263 nm and 240 nm get changed in the presence of compounds. (II) Showing formation of mixed hybrid G-quadruplex structure in TelQ in the presence of compounds A-D. The peak intensity gets changed in the presence of compounds

### 3.2 Thermal stabilization of G-quadruplex

The CD melting method was used to investigate the thermal stabilization of both of the chosen G-quadruplex oligos in the presence of compound A-D. Plotting the fraction-folded versus temperature graph allowed for the calculation and comparison of each transition’s melting temperature (Tm). The Tm of PRG4Q was recorded at approx 60°C which get increases to 66.5°C, 64°C, 66.9°C, and 67°C in the presence of compound A-D respectively. Similarly, Tm of TelQ was found to increase from 58.7°C to 61.47°C, 63.5°C, 63.2°C, and 63.9°C (Figure 4).

**Figure 4:**
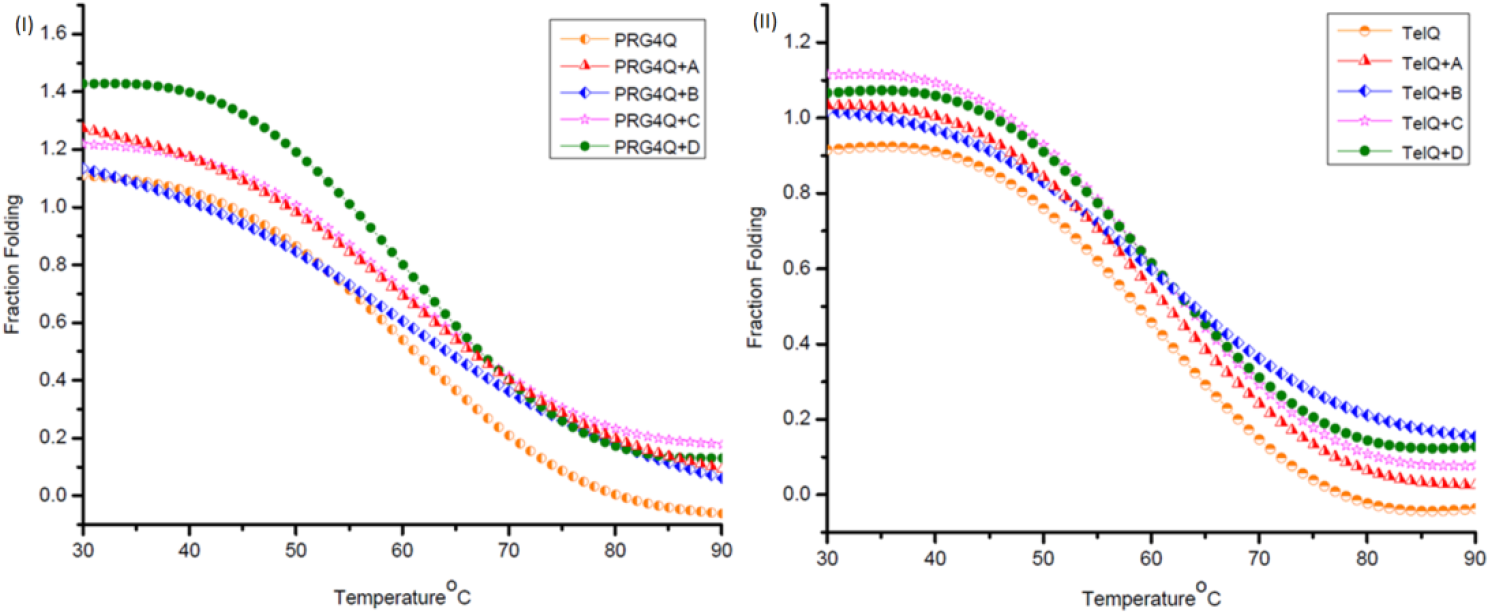
CD melting curve of 5 μM PRG4Q and TelQ in the absence or presence of 10 μM compounds A-D. (I) Showing increment in melting temperature (Tm) of PRG4Q in the presence of compounds A-D. (II) Showing increment in melting temperature (Tm) of TelQ in the presence of compounds A-D

### 3.3 UV-Vis Spectroscopy

Compound A-D’s interactions with both G4Q structures were examined using UV-Vis absorption spectroscopy. Table 1 lists the compounds’ maximum absorption wavelength. Figures 5 and 6 illustrate that as the concentration of G4Q oligos is increased from 1 to 10 μM, the absorption peak of each compound decreases, indicating the development of compound-G4Q or an interaction between the compound and G4Q. On the other hand, no broad change in peak intensity was found with the addition of G4M, mutated G4Q sample (Figure 7) as well as dsDNA (Figure 8) with any of the 4 compounds. These results support the selective interaction of each of the selected compounds (A-D) with of G-quadruplex conformation rather than non G-quadruplex or duplex conformation.

**Figure 5:**
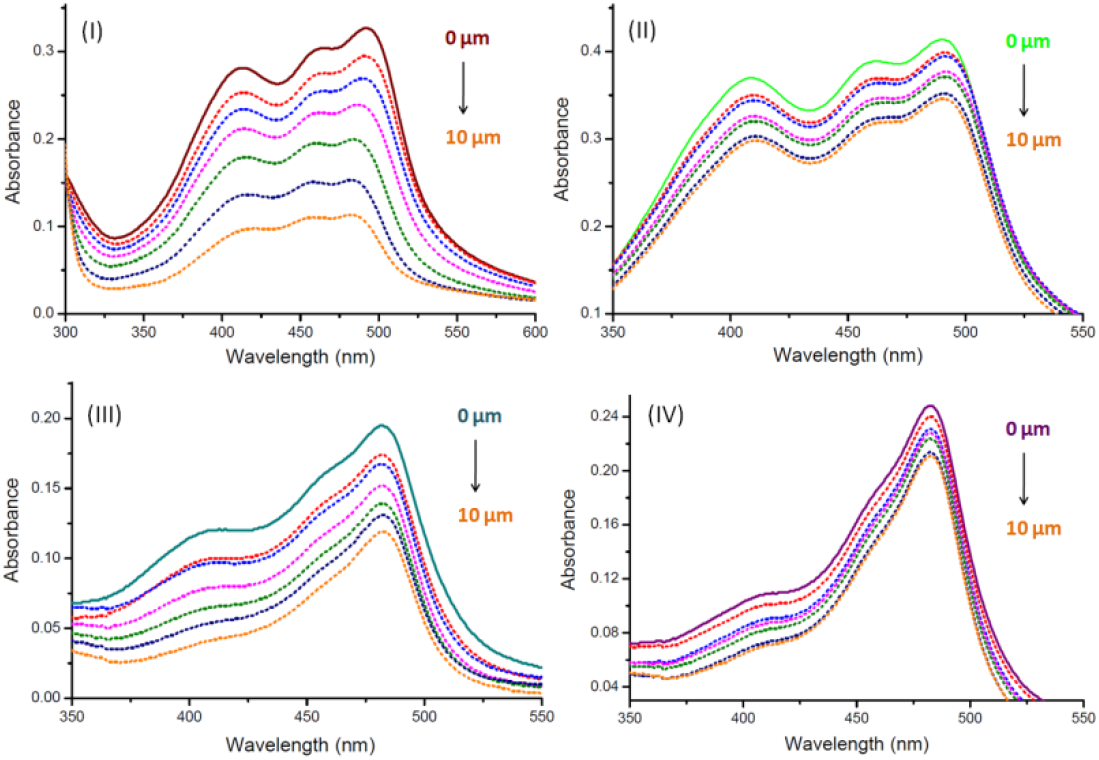
Absorption spectra of compound A-D in the absence and presence of increasing concentration (0,1, 2, 4, 6, 8, and 10 μM) of PRG4Q. Absorption peaks of compound A (I), B (II), C (III) and D (IV) get decreases as PRG4Q’s concentration increases.

**Figure 6:**
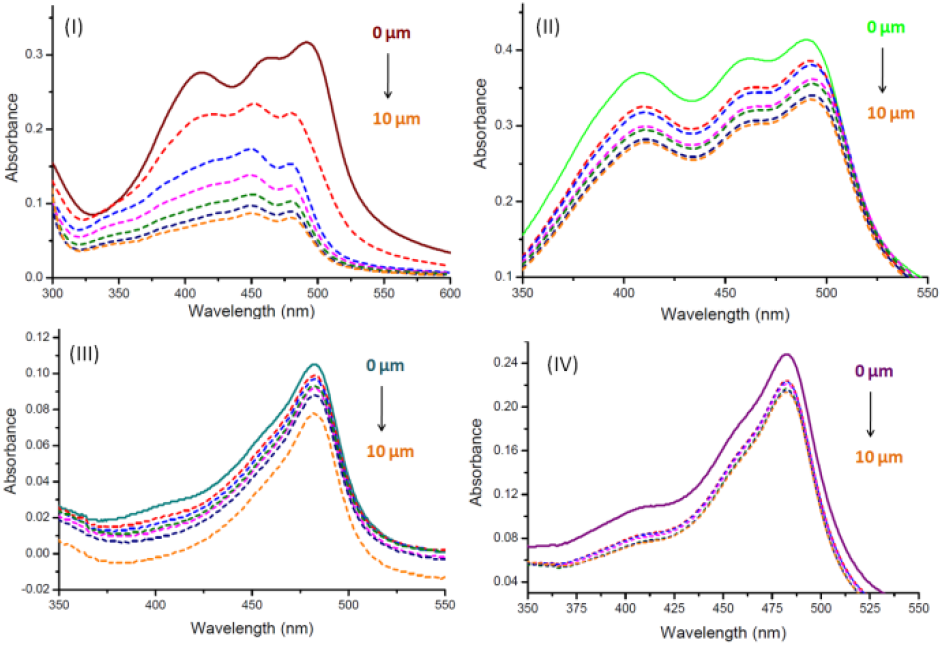
Absorption spectra of compound A-D in the absence and presence of increasing concentration (0,1, 2, 4, 6, 8, and 10 μM) of TelQ. Absorption peaks of compound A (I), B (II), C (III) and D (IV) get decreases as TelQ’s concentration increases

**Figure 7:**
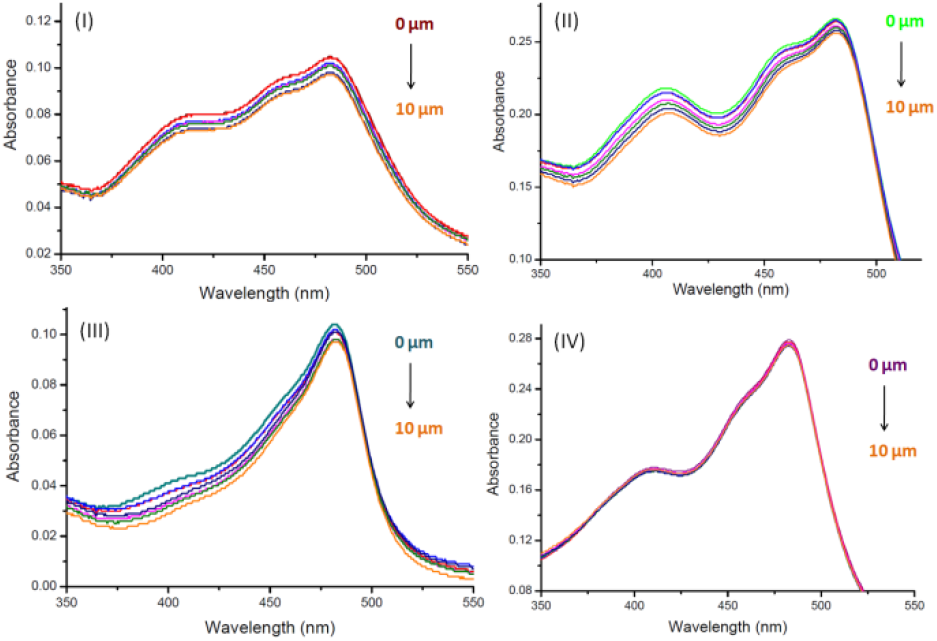
Absorption spectra of compound A-D in the absence and presence of increasing concentration (0,1, 2, 4, 6, 8, and 10 μM) of G4M. Absorption peak of compound A (I), B (II), C (III), and D (IV) does not get changed with increasing concentration of G4M

**Figure 8:**
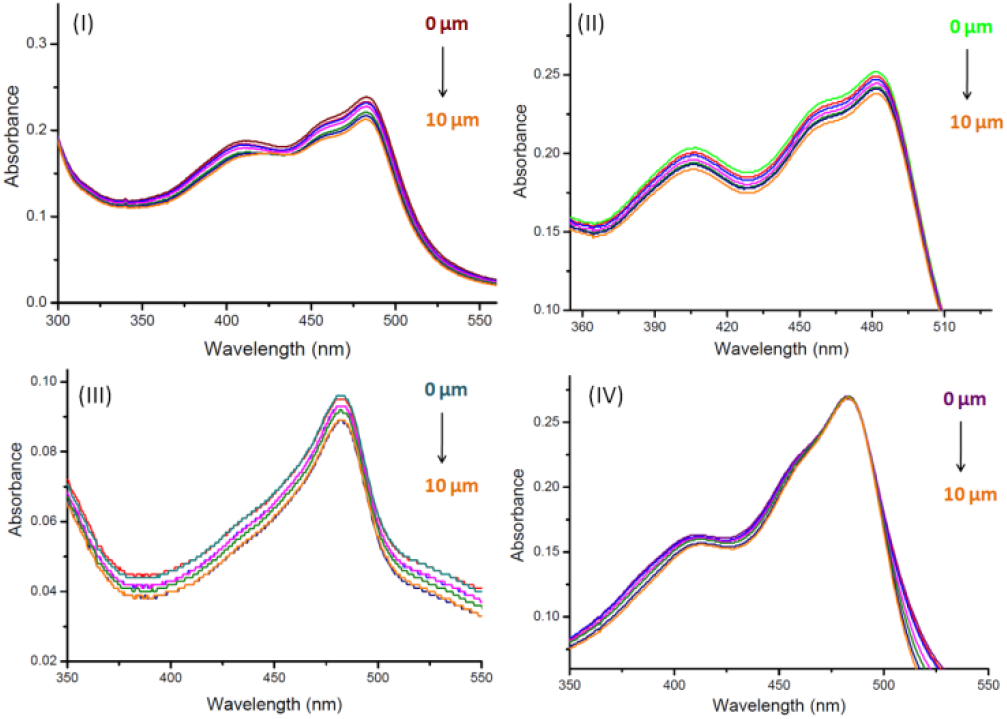
Absorption spectra of compound A-D in the absence and presence of increasing concentration (0,1, 2, 4, 6, 8, and 10 μM) of salmon sperm duplex DNA (dsDNA). Absorption peak of compound A (I), B (II), C (III), and D (IV) does not get changed with increasing concentration of dsDNA

### 3.4 Ligand binding study with fluorescence spectroscopy

Fluorescence enhancement and quenching in compound A-D was observed during the interaction of folded G-quadruplex samples with compounds. Excitation and emission maxima of each compound are mentioned in Table 1. With PRG4Q, chroman ligand A and its derivative complex B causes fluorescence to be enhanced, whereas chroman ligand C causes fluorescence to be quenched, and its derivative complex D cause fluorescence to be enhanced (Figure 9). However, ligand A and C both exhibit fluorescence quenching in response to TelQ, whereas their complexes B, and D both exhibit fluorescence enhancement in response to TelQ (Figure 10). Additionally, there was not a noticeable variance in the fluorescence intensity of any compound in the G4M and dsDNA samples (Figure 11-12), indicating that there was no interaction between the DNA and compounds. This is in support of the compounds’ preference for binding to G-quadruplex conformation over duplex or other conformations of DNA.

**Figure 9:**
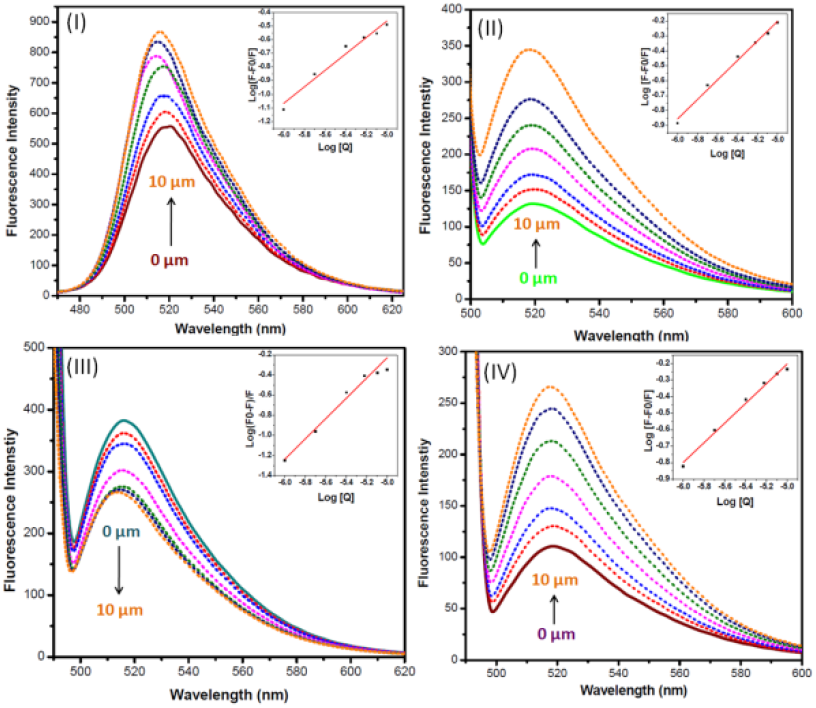
Fluorescence emission spectra of 30 μM compound A-D in the presence of increasing concentrations of PRG4Q (0, 1, 2, 4, 6, 8, and 10 μM), Emission peak of compound A (I), B (II), C (III), and D (IV) gets changed with increasing concentration of PRG4Q. The double logarithmic plot to determine Ksv and K is shown in the inset

**Figure 10:**
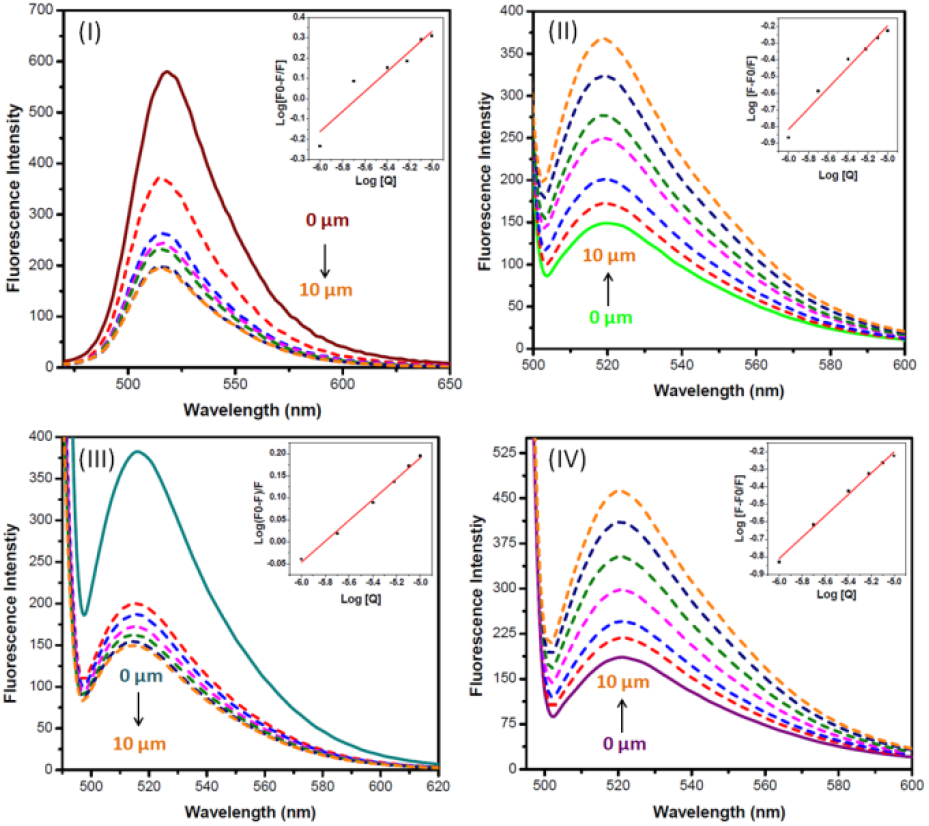
Fluorescence emission spectra of 30 μM of compound A-D in the presence of increasing concentrations of TelQ (0, 1, 2, 4, 6, 8, and 10 μM), Emission peak of compound A (I), B (II), C (III), and D (IV) gets changed with increasing concentration of TelQ. The double logarithmic plot to determine Ksv and K is shown in the inset

**Figure 11:**
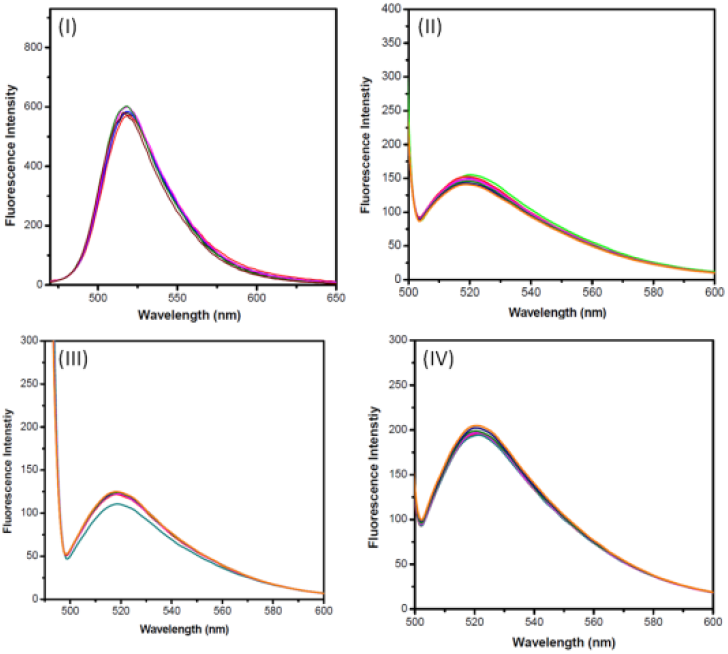
Fluorescence emission spectra of 30 μM of compound A-D in the absence and in the presence of increasing concentration (0,1, 2, 4, 6, 8, and 10 μM) of G4M. Emission peak of compound A (I), B (II), C (III), and D (IV) does not get changed with increasing concentration of G4M.

**Figure 12:**
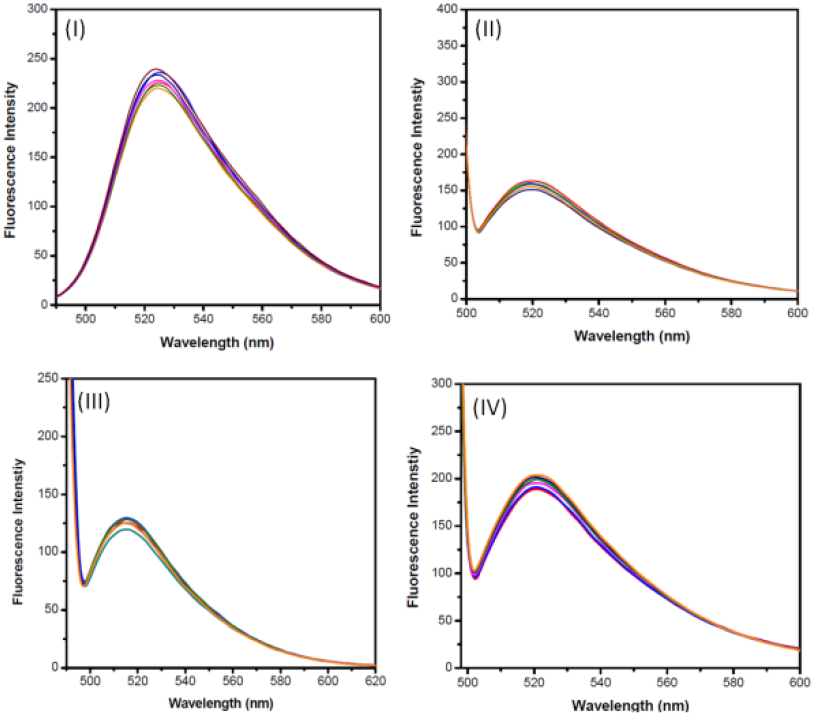
Fluorescence emission spectra of 30 μM of compound A-D in the absence and in the presence of increasing concentration (0,1, 2, 4, 6, 8, and 10 μM) of salmon sperm duplex DNA (dsDNA). Emission peak of compound A (I), B (II), C (III), and D (IV) does not get changed with increasing concentration of dsDNA

Further using the Stern-Volmer equation (Eq.1) (Lakowicz, 2006; Phopin et al., 2019), fluorescence data were analyzed and double log plots of F_0_-F/F versus [Q] for fluorescence quenching and F-F_0_/F versus [Q] for fluorescence enhancement were generated.

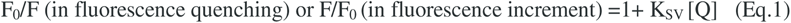

Where Q is ligand concentration, Ksv is the Stern Volmer constant, and F and F_0_ are the fluorescence intensities of compounds in the presence and absence of the G4Q DNA samples, respectively. To assess binding affinity the modified Stern-Volmer equation (Eq.2 and 3) was used (Phopin et al., 2019).

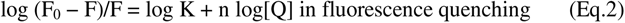

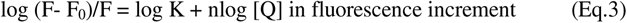

K and n are the corresponding binding constant and number of binding sites. The Stern Volmer constant (Ksv) and the binding constant (K) of each compound with both G-quadruplex samples were listed in Table 2. The values of K for the G quadruplex-compound interaction were calculated from the intercept and slope of the plot of log (F_0_-F)/F or log (F-F_0_)/F vs. log [Q] (Figure 9-10).

**Table 2:**
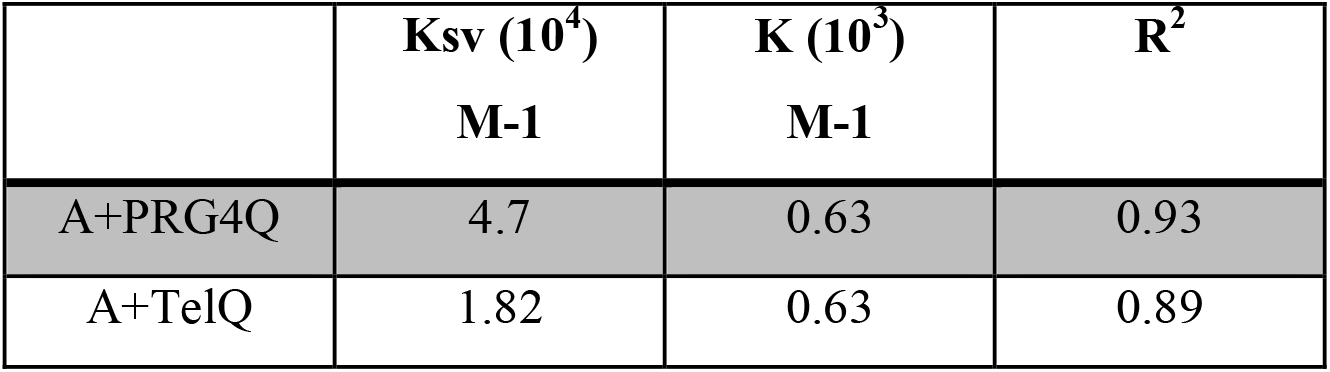
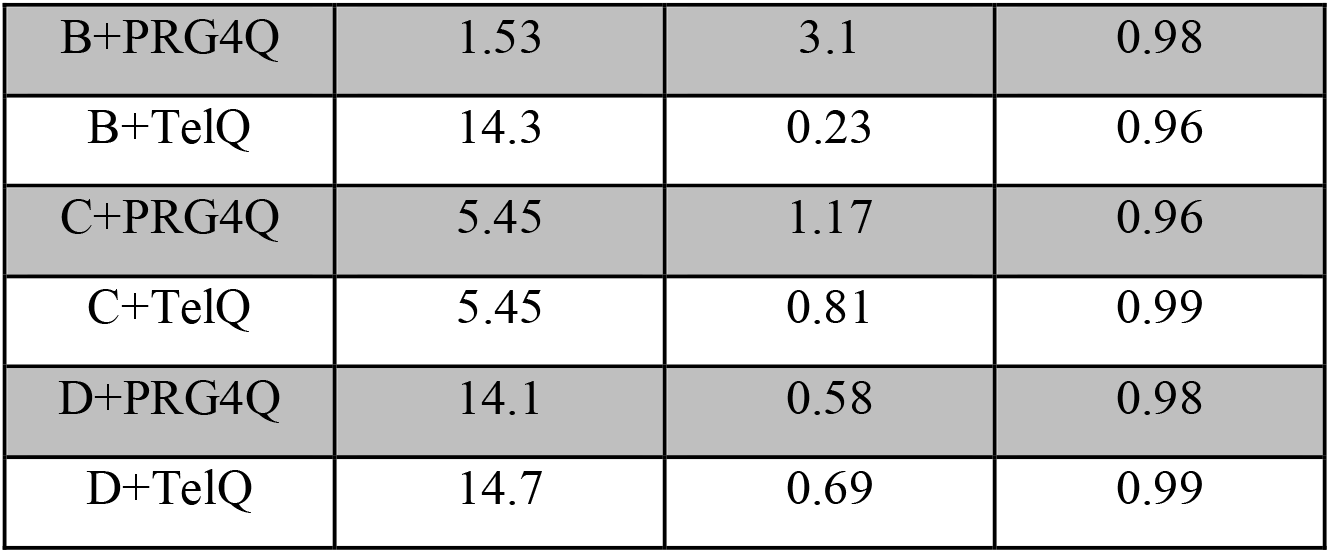
Values of Ksv and binding constant (K) of compound A-D with PRG4Q and TelQ. R^2^ is correlation coefficient

### 3.5 Fluorescence Intercalating Displacement assay

FID curves were created by graphing a percentage of TO displacement versus the concentration of particular compounds (Figure 13). Using the formula PD = 100 - [(FA/FA_0_) 100], the fluorescence area (FA) was transformed to percentage displacement (PD). FA_0_ and FA are the Fluorescence area without or with any chemicals added respectively. By using a linear regression fitting curve, the concentrations of chemicals at which 50% displacement of TO (DC50) occurs were determined from the FID plot. As for the PRG4Q, a concentration of 30, 36, 33, and 25.7 μM of compounds A, B, C, and D respectively were found to require displacing 50% TO, indicating a strong affinity of these compounds to parallel G4Q topology. On the other hand, DC50 values for TelQ were found at concentrations of 47, 36.8, 42, and 33.8 μM for compounds A, B, C, and D respectively suggesting strong affinity of both complexes B, and D for TelQ compared to their chroman ligands.

**Figure 13:**
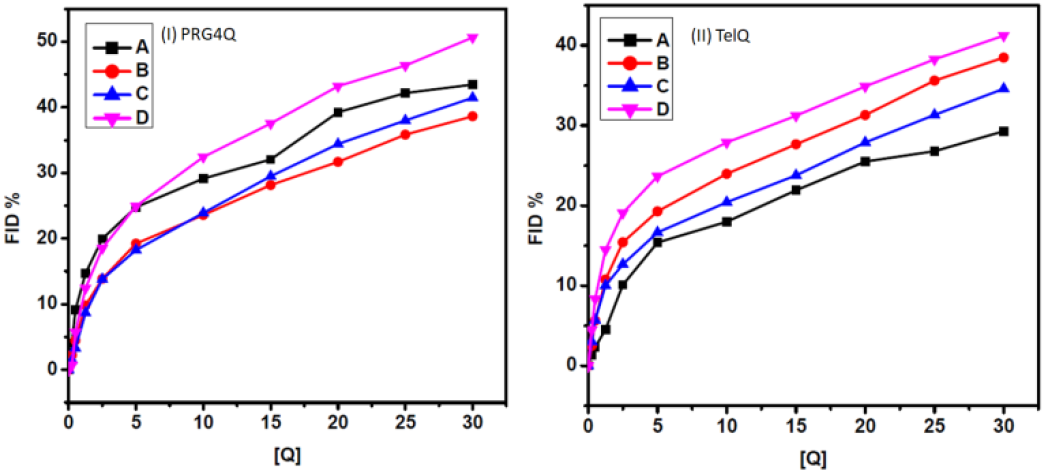
FID curve of compound A-D with increasing concentration of PRG4Q (I), and TelQ (II) obtained by plotting %FID vs. concentration.

### 3.6 Cytotoxicity

Given their utility as G-quadruplex targeting ligands in a cellular context, it is intriguing to explore the toxicity of these chroman ligands and their complexes towards HEK293T cells. For this MTT assay was used to evaluate the toxicity of these compounds on HEK293T cells. As compounds were found to crystallize in media at concentrations above 50 μM, cells were treated with compounds at increasing concentrations up to 50 μM as a maximum concentration. Figure 14 depicts a rise in cell death percentage with compound concentration, although the maximum cell death % at 50 μM concentration was only determined to be between 35 to 45%. The IC50 concentrations for ligands A and C as well as complexes B and D were 77, 58, 57, and 69 μM, respectively.

**Figure 14:**
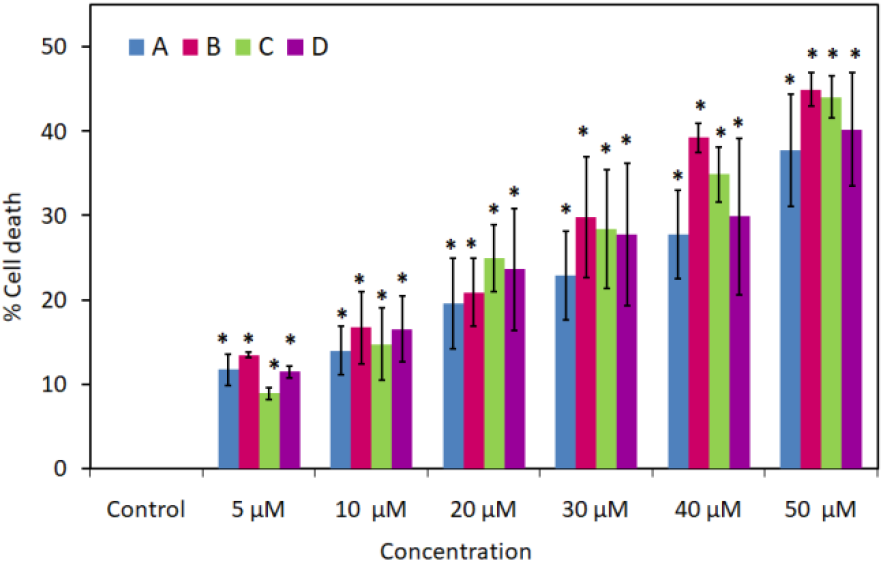
MTT graph showing antiproliferative effect of compounds A-D on HEK293T cell line. The error bar shows the mean and SD. The paired t-test was used to measure statistical significance. *P ≤ 0.05 compared with control vs. treatment groups

### 3.7 Cellular imaging of G-quadruplex

As seen in Figure 15, HEK293T cells treated with 20 μm of each compound individually increased the signal of BG4 or G-quadruplex foci as compared to untreated cells. The presence of BG4 signal or G-quadruplex foci inside the nucleus and around the cytoplasm, which stand for DNA and RNA G-quadruplexes, respectively, was observed (Biffi et al., 2013, 2014). Compound-treated cells exhibit considerably higher levels of both of these foci. Endogenous BG4 signals had an average fluorescence intensity of 14.39 in control cells, whereas this increased to 28.45, 25.09, 22.96, and 23.82 in compound A-D respectively treated cells.

**Figure 15:**
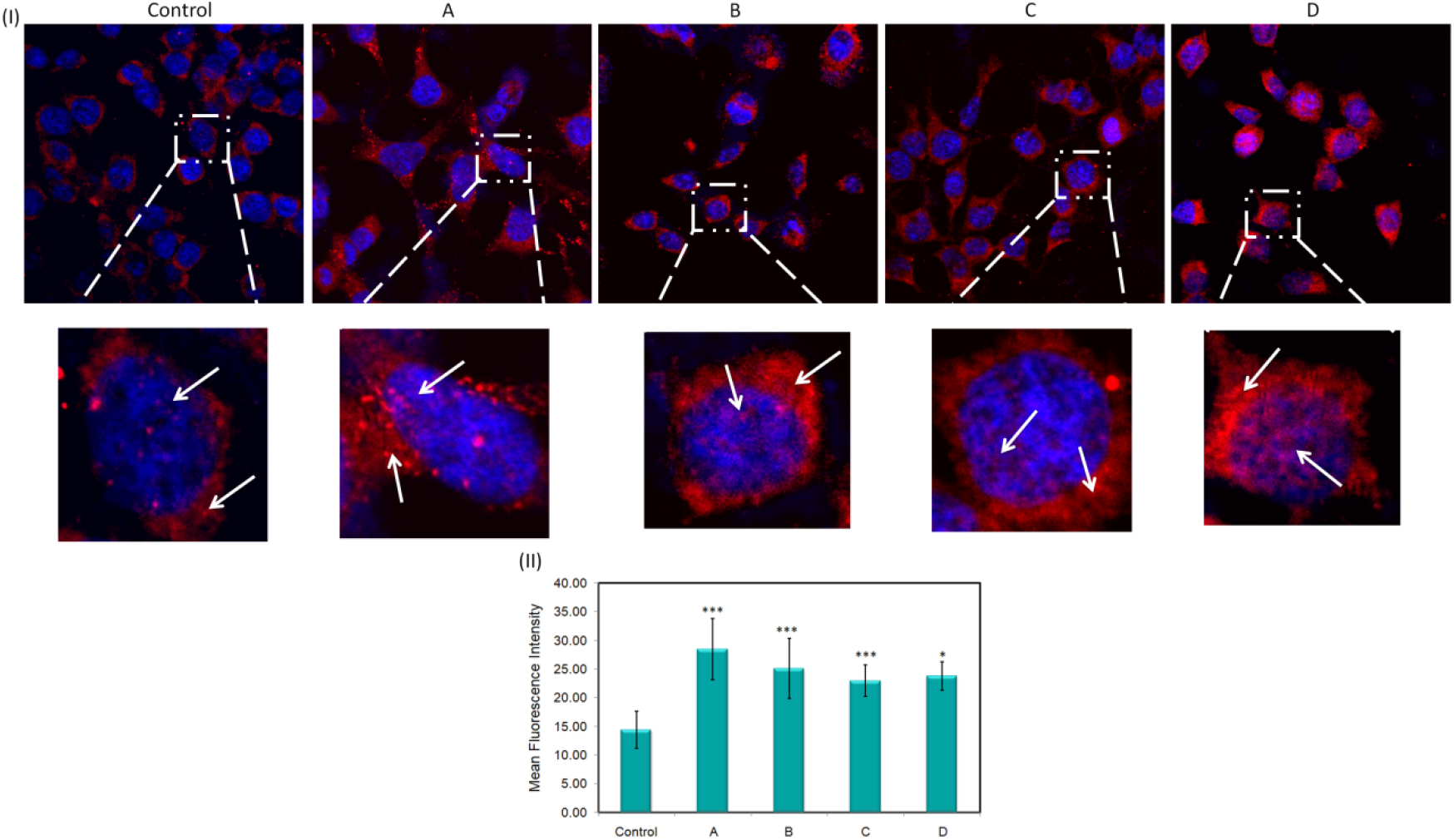
(I) Confocal imaging of G-quadruplex foci in HEK293T cell with or without compound A-D treatment. G-quadruplex foci (red) has been enhanced in cells with compound treatment, suggests increment in G-quadruplex formation compared to without treatment. Nuclei were counter stained by DAPI. Scale bar 33 μm. (II) Bar graph of mean fluorescence intensity of endogenous G-quadruplex foci. Error bars represent the S.D. of three independent experiments. *P ≤0.05, without compound vs. with compound

### 3.8 Molecular docking and binding profile

The binding scores of the parallel G-quadruplex that were docked with the ligands A, C and complexes B, and D were 3516, 3732, 3314, and 3812, respectively. However, when docked with human telomeric DNA G-quadruplex, revealed binding scores of 3370, 3568, 3724, and 4182 for ligands A, C and complexes B, and D respectively. The non covalent interactions of docked complex showed in Figure 16 and supplementary file (S1). Hydrogen and hydrophobic bonding, as well as pie (π) stacking, are observed in the parallel G-quadruplex with ligand A, hydrophobic and π-stacking with ligand C and complex D, and hydrogen bonding and π-stacking with complex B. Mixed hybrid G-quadruplex, on the other hand, exhibits hydrogen bonds with ligand A, hydrogen and hydrophobic bonds, and π-stacking with ligand C, hydrogen bonds, water and salt bridges, metal complexes, and π-stacking with complex B, and hydrogen bond and π-stacking with complex D. These interaction data point to π - π interactions between chosen chroman ligands and their complexes with G-quadruplex. Parallel G-quadruplex interactions with these chroman ligands and complexes were found to be end-stacked, whereas interactions with mixed hybrid quadruplex structures were groove-bound (Figure 17).

**Figure 16:**
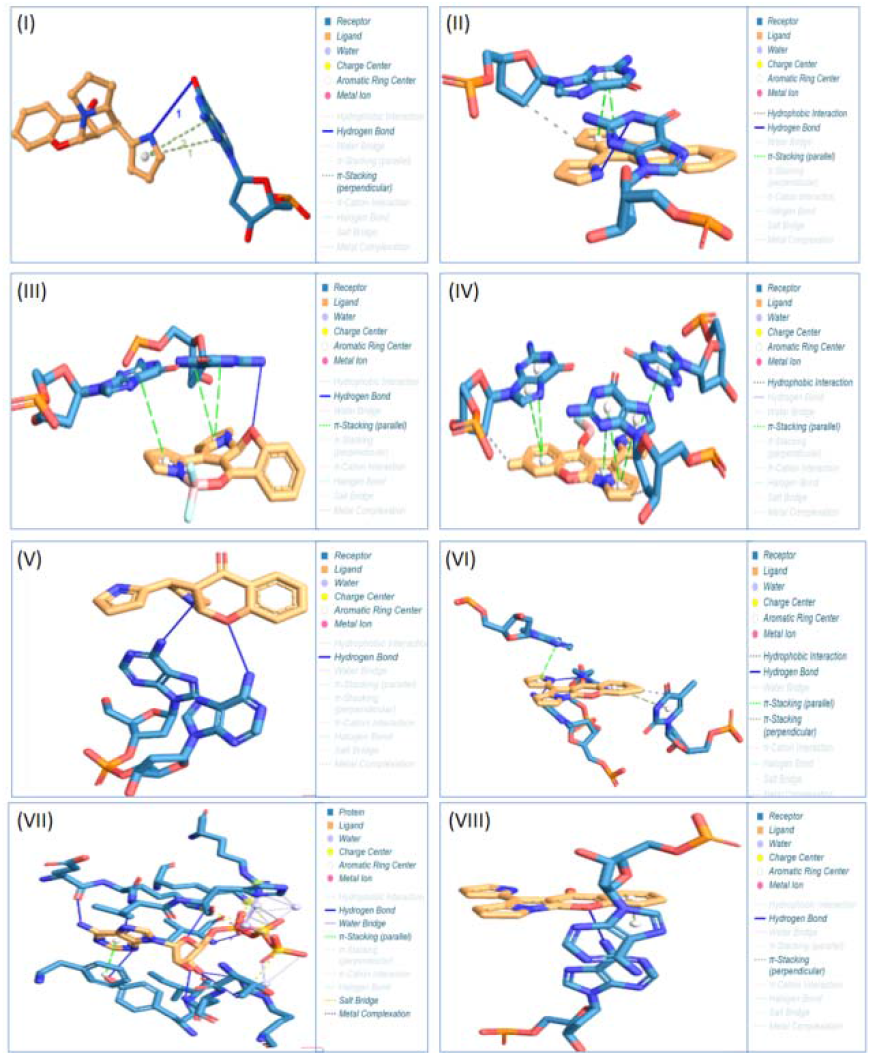
Schematic presentation of docking showing annotation of non-covalent interactions of compounds A to D with parallel and hybrid-type telomeric G-quadruplexes obtained with Patch Dock. Parallel G-quadruplex with Ligand A (I), ligand C (II), complex B (III), complex D (IV). Mixed hybrid-type G-quadruplex with ligand A (V), ligand C (VI), complex B (VII), and complex D (VIII)

**Figure 17:**
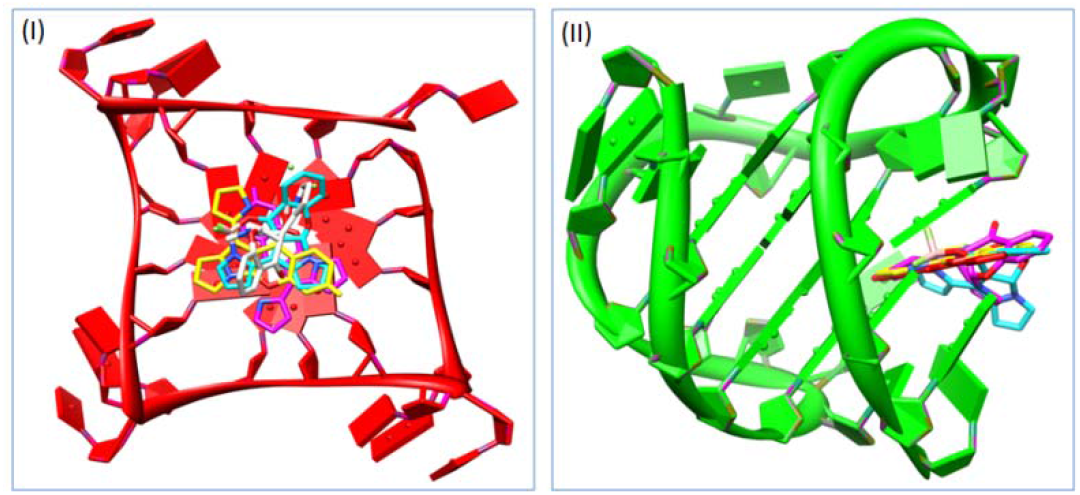
Schematic 3D presentation of the interaction pattern of the DNA-Ligand complex (I) parallel G-quadruplex interaction with all four compounds (A-D), showing end-stacked interaction. (II) Mixed hybrid-type G-quadruplex interaction with all four compounds (A-D), showing groove binding patterns.

## 4. Discussion

In this study, we focused on to demonstrate the G-quadruplex binding and stabilizing by the interaction of highly fluorescent chroman derivatives (A and C) and their corresponding borondifluoride complexes (B and D) which was synthesized as a meso-substituent based on dipyrrins. Compounds A-D was preliminarily screened for their ability to stabilize G4Qs by using a circular dichroism (CD) melting assay. Several diverse G4Q-forming sequences able to form parallel, antiparallel, and hybrid G-quadruplex structures. In this study two G4Q-forming oligonucleotides were used. One is PRCC G4Q, 27 bp long sequences within exon 1 of *PRCC* which also exist in 5’ end of *PRCC-TFE3* fusion gene (Das & Verma, 2020; Neha & Das, 2023) and another is 25 bp long human telomeric sequences. In order to establish the folding of each G-quadruplex structure in both oligo samples, CD spectra were first obtained. CD spectra of folded oligo samples show that parallel G-quadruplex structures are formed in PRCC G4Q while mixed hybrid conformations are formed in human telomeric sequences. Additional CD experiments were performed to verify the capability of compounds A-D to alter the native folding topology of these G4Qs. No relevant differences in the CD profiles were detected for any of the analyzed G4Qs however peak intensity get changed in the presence of each individual compound, suggesting a general preservation of each G4Q architecture upon compound addition. The stabilizing properties of A-D for G-quadruplex structure were then evaluated by CD-melting experiments by measuring the compound-induced change in the melting temperature (ΔTm) of G4Qs. The melting temperature (Tm), where the complex is 50% dissociated, represents the midpoint of a particular transition. The Tm value basically gives a general evaluation of the prevailing form. The structure is mostly folded at temperatures below the Tm and mostly unfolded at temperatures above the Tm (Mergny & Lacroix, 2009). DNA melting experiment showed an enhancement (approx 4-7°C) in Tm of G4Q samples with the addition of each individual compound which clearly indicates a good G4Q-stabilizing effect for all the investigated compounds, In addition, the highest thermal stabilization effects were observed in complex D for both G4Qs topologies. A ligand’s DNA binding behavior is routinely studied using UV-visible absorption. In this work, we examine how each of the four chemicals (A-D) binds to the G-quadruplex. When compared to duplex or non-G-quadruplex structure, UV-Vis spectroscopy showed that all four chemicals interact preferentially with the G-quadruplex structure. The hypochromism without significant changes in spectral wavelength and absence of isosbestic point was observed in the absorption spectrum. In contrast to G4M and duplex DNA, G-quadruplex causes a greater variation in hypochromicity in compound A-D (30-60%) with no change in the compounds’ Soret band wavelength. The binding interaction of substances with the G-quadruplex is established by this hyporchromism without changing peak position. We also looked into whether a compound’s fluorescence characteristics were impacted by G-quadruplex binding. Since each emissive species in solution contributes to the steady state fluorescence spectrum, it may be used to track changes in the chromophore’s surroundings. Fluorescence quenching is the term for the fall in a fluorophore’s quantum yield resulting from a variety of chemical bonding with a quencher molecule, such as excited-state reactions, molecule rearrangements, energy transfer, the creation of ground state complexes, and collisional quenching. Static and dynamic quenching are two common forms of quenching. By colliding with the quencher, the fluorescent material causes dynamic quenching, which results in a reduction in quantum yield. The absorption spectra of a fluorescence material are unaffected by dynamic quenching, but they are affected by static quenching because a compound is created between the ground state of the fluorescence substance and the quencher (Lakowicz, 2006; Pan et al., 2011; Papadopoulou et al., 2005). Fluorescence quenching and enhancement cannot happen in either scenario without their being molecular interaction between the fluorophore and the quencher. The accessibility of the fluorophores to quenchers can also be revealed by using the fluorescence quenching method (Papadopoulou et al., 2005). Fluorescence spectrum of chroman derivative ligand A with PRG4Q and TelQ found to increase and quenched respectively, while ligand C results in quenching with both PRG4Q and TelQ. On the other hand both borondifluoride complexs B, and D with both PRG4Q and TelQ shows fluorescence enhancement. The fall and increment in compounds’ fluorescence intensity suggest that the substance’s ability to bind to the G-quadruplex has effectively declined and increased in quantum efficiency respectively (Lakowicz, 2006; Papadopoulou et al., 2005). These changes in the fluorescence intensities of both ligands and their complexes were used to determine binding constant. Additionally, Ksv (Stern-Volmer constant) was calculated because it is regarded as a measure of the effectiveness of fluorescence quenching or increment by DNA. All four compounds’ Ksv values were found to be high, pointing to the intercalative binding mode of compounds with G-quadruplex topology. Furthermore, Ksv and binding constant values show that ligands C and its complex D strongly interact with both the parallel (PRG4Q) and mixed hybrid G-quadruplex (TelQ) topologies, whereas ligand A and complex B strongly interact with the parallel and mixed hybrid topology respectively. It appears that ligand A and compound B have a modest affinity for mixed hybrid and parallel configuration, respectively. The enhancement of fluorescence intensity of compounds A, B, and D on binding with PRG4Q and TelQ accordingly is due to an increase in the quantum yield of compounds upon binding with G-quadruplex, which suggests external groove binding as the mechanism of this interaction. While enhanced fluorescence was previously thought to be connected with solely intercalative binding, these types of binding cause a slight or no spectrum change with an occasional hyperchromicity (Sari et al., 1990; Sehlstedt et al., 1994). Later research has demonstrated its occurrence both during groove binding and G-quadruplex-ligand interaction. A further finding that supports the selective binding of compounds with G-quadruplex conformation rather than a duplex or non-G-quadruplex conformation was the absence of any significant change in the fluorescence intensity of any molecule in G4M and dsDNA samples, indicating no interaction between the DNA and compounds. FID studies were conducted to gain insights into compound A-D’s affinity for G4Qs. Briefly, this test is based on the displacement of a fluorescent dye, thiazole orange (TO), from DNA following the addition of increasing concentrations of a candidate molecule. The FID assay is a straightforward and efficient approach for determining the affinity of different ligands for binding quadruplexes. The competitive displacement of the bound probe by ligands enables the quantification of the binding constant for substances (Monchaud et al., 2008). With a high affinity, the dye thiazole orange end-stacks on the two exterior quartets of a G-quadruplex. Because TO is entirely nonfluorescent when free in solution but extremely fluorescent when interacting with G-quadruplex DNA, the displacement of a particular ligand can be easily tracked by a decrease in TO’s fluorescence (Tran et al., 2011). The ligand-induces TO displacement was quantified using the DC50 value, which was used to determine the ligand concentration required to remove 50% of the TO from G-quadruplex DNA. The ligands were then sorted using this value. Because compounds B, C, and D for TelQ and A, C, and D for PRG4Q, respectively, dissociate 50% TO at lower concentrations than A and B in TelQ and PRG4Q, respectively, DC50 values show that ligands A for TelQ and complex B for PRG4Q have weak affinities in comparison to other compounds. This is also supported by Tm and fluorescence binding analysis data. All of these biophysical studies on ligand-DNA binding point to chroman ligand C and its borondifluoride complexe D have a higher propensity for binding with G-quadruplex than A and B. To determine the binding poses and relative binding affinities of all four compounds with both G-quadruplex topologies, a molecular docking study was carried out. We selected the parallel conformation of the human telomeric G-quadruplex for the docking analysis because the crystal structure PRCC G-quadruplex is not accessible in the PDB database. Interestingly binding score between the ligand and complexes with both G-quadruplex topologies supports our biophysical findings and suggests the interaction between both ligand and complexes with G-quadruplex structure. The *In silico* data also indicate that, out of the four compounds examined, complex D is interacting with the G-quadruplex the most as it showing highest binding score with both G-quadruplex topologies. Non covalent interaction of docked complex suggests π-π interaction which can contribute to the binding affinity and specificity. They enhance the stability of the complex by providing additional non-covalent interactions.The binding affinity, selectivity, and structural impacts of the interacting complexes are believed to be improved by hydrogen bonding, water and salt bridges, and hydrophobic interactions. The overall binding strength is influenced by these non-covalent interactions (Bosshard et al., 2004; Neidle & Sanderson, 2021; Ricci & Netz, 2009; Spyrakis et al., 2007). Based on our *In vitro* biophysical and *In silico* interaction data, it can be said that these ligands and their complexes successfully interact with the G-quadruplex structure by end stacking mode with parallel and external groove binding pattern with mixed hybrid G-quadruplex topologies. End stacking occurs when a ligand molecule attaches to the quadruplex structure’s exposed ends, often at the terminal G-quartets. The aromatic rings of the ligand align and stack on top of the aromatic rings of the guanine bases, facilitating π-π stacking interactions. A hybrid quadruplex’s ligand molecules interact with the groove, which implies that they bind to and interact with the unique structural elements of the quadruplex DNA molecule. Multiple interaction patterns can be at play when molecules interact with the groove of a hybrid quadruplex. The chemistry of the ligand molecule and the quadruplex’s structural features determine the precise interaction pattern. To gain insight into the versatility of these ligands and their complexes for the G-quadruplex detection or stabilization inside the cells, Confocal imaging of cells incubated with these compounds was done. Since a gene’s capacity to form G-quadruplexes is regulated by the progression of the cell cycle and endogenous G-quadruplex structures can be stabilized by a small-molecule ligand or G-quadruplex stabilizer, HEK293T cells were individually treated with 20 μM of each of these four compounds for 24 hours. Because 20 μM is found to be non-toxic to cells according to MTT data, this concentration is used in the cellular experiments. Comparing cells treated with compounds to cells not treated, a substantial increase in BG4 signal was observed. Therefore, we expected that giving cells a compound treatment would make them good G-quadruplex targets for the BG4 antibody. To track the G-quadruplex development within the cells, fluorescence intensity was monitored and compared. This G-quadruplex-specific assay used the G-quadruplex-specific antibody BG4, which can recognize G-quadruplex in human cells’ DNA (Biffi et al., 2013) and RNA (Biffi et al., 2014) strands. Due to the FLAG tag sequence on the BG4 antibody, it can be recognized by a FLAG antibody utilizing an anti-Flag Alexa Fluor 546 secondary antibody. The average fluorescence intensity of antibodies bound to G-quadruplexes was found to be higher in compound-treated cells than in untreated control cells. Numerous PQS-containing genes, including telomeres, are present in millions of human cells. These PQS have the ability to fold into a G-quadruplex structure during the S-phase or DNA replication phase (Biffi et al., 2013), which may help to explain why signals were seen in control cells. Together, these findings show that these recommended compounds interact with and stabilize the G-quadruplex structure. This interaction and stabilization are also maintained in cellular environments, which may open up opportunities for investigating G-quadruplex target studies in both *In vitro* and cellular contexts. However, some limitations exist in terms of specificity; the strong fluorescence of these compounds prevents them from being used for precise labeling of G-quadruplex in cells because cells were observed to totally stain after being incubated with the compounds (unpublished data). To get over this restriction and create a new generation G-quadruplex marker that can be used in ongoing cellular G-quadruplex imaging research, we will aim to find applications in chemical biology quests.

## 5. Conclusion

In conclusion, we looked at the G-quadruplex binding properties of a few highly fluorescent chroman-BF2 complexes and investigated the influence of substituents and the chemical nature of the interaction. The production of G-quadruplex structures with the right ligand is a crucial goal in the creation of new drugs. Our research demonstrates that these chromam ligands and their complexes have the ability to bind and stabilize G-quadruplex topologies in parallel and mixed hybrid forms both *In vitro* and within living cells. Our research as a whole point to these chroman complexes as a potentially useful chemical compound for G-quadruplex-specific ligands and broadens the range of potential G-quadruplex targeting ligands.

## Acknowledgement

We acknowledge Central Discovery Centre (CDC), Banaras Hindu University, Varanasi for Circular Dichroism facility. We acknowledge Dr. Gopeshwar Narayan, Professor, Molecular and Human Genetics, Banaras Hindu University for providing facility of microplate reader. We acknowledge ISLS, Banaras Hindu University for providing facility for confocal microscopy.

